# The Energetic Cost of Adrenergic Signaling in Primary Human Fibroblasts

**DOI:** 10.64898/2026.07.09.737569

**Authors:** Janell L.M. Goode, Gabriel Sturm, Martin Picard

## Abstract

Stress involves the activation of cellular, physiological, and emotional processes that cost energy—nothing is free in biology. In mammals, the stress response involves hormone release, including norepinephrine (NE), which increases energy expenditure. To quantify the energetic cost of NE signaling in a simple cellular system, we interrogated the dose (0 – 10 μM NE) and time-dependent (up to 10 hours) effects of adrenergic signaling in primary human fibroblasts. Oxygen consumption rates (OCR, reflecting ATP generated by mitochondria) and extracellular acidification rate (ECAR, reflecting ATP generated by glycolysis) were measured continuously using extracellular flux analysis, allowing us to estimate the ATP turnover rates, and thus cellular energy expenditure. Within the first 18 minutes (early phase), glycolysis increases up to 47% whereas respiration decreased 2-5%. Both parameters normalized within 1-2 hours for low NE concentrations. This was followed by an increase in oxidative phosphorylation (OxPhos), peaking around 9-12% by 2-6 hours (mid or late-phase). These minutes-to-hours data reveal the temporal dynamics whereby NE increases cellular energy expenditure in fibroblasts. Blocking OxPhos with oligomycin or piericidin A abolished OxPhos changes post-NE addition while conserving the glycolytic response. Withdrawal of glucose from the media significantly dampened the absolute rise in ECAR in response to NE, and instead increased OxPhos, revealing the metabolic flexibility in fibroblasts. Finally, cells with genetic defects impairing OxPhos exhibited a 50% blunted NE-driven metabolic response, consistent with the existence of an energy constraint in mitochondrial diseases. In summary, we have resolved the dynamics and flexible bioenergetic recalibrations associated with NE-driven hypermetabolism in primary human fibroblasts. Mapping the nature and magnitude of these recalibrations in humans would advance our understanding of the potential energetic forces underlying the damage to health by chronic stress.

## Background

Stimuli that threaten homeostasis^1^ cause energy-dependent stress responses across the organism. As a result, the brain activates the sympathetic-adreno-medullar (SAM) axis, which releases catecholamines such as norepinephrine (NE) from sympathetic nerve terminals as well as the adrenal medulla^2^. NE functions both as a neurotransmitter within the central nervous system and as a hormone peripherally^2^. NE acts on adrenergic receptors, a family of g-coupled protein receptors (GPCRs), which when activated start a cascade of second messengers within the cell^3^. This leads to organ wide responses such as increased heart rate, the redirection of blood flow to necessary organs, and promote substrate mobilization from energy stores^2^. At the organismal level, the systemic metabolic effects of catecholamines are well characterized.

Activation of the SAM axis elicits a coordinated systemic response, including increased heart rate, redirecting blood flow to essential and away from non-essential organs, and the mobilization of substates such as glucose and fatty acids from energy stores^2,4^.

At the cellular level, energy transformation can occur via two principal pathways: oxidative phosphorylation (OxPhos), which generates ATP in the inner membrane of mitochondria^5^, and glycolysis, which generates ATP through the conversion of glucose to pyruvate in the cytoplasm^6^. Catecholamines are well-established regulators of systemic energy expenditure, yet the cell-autonomous bioenergetic consequences of adrenergic signaling is still underway. In particular, how norepinephrine causes individual cells to recalibrate their energy transforming pathways remains completely understood.

To address this question, we use primary human fibroblasts, a cellular model expressing adrenergic receptor subtypes^7,8^ widely used for mitochondrial and cell metabolism studies^9,10^. We previously used human fibroblasts to explore the bioenergetic costs of chronic glucocorticoid signaling^10^. The cells recalibrated their adrenergic receptor expression in response to chronic glucocorticoid activation^8^. They also offer a relatively easy means of obtaining patient derived samples, such as those from mitochondrial disease patients with mutations in SURF1, an assembly factor for complex IV of the electron transport chain, that can then be used to dissect questions aimed at when one ATP producing pathway is deficient^9,11^. Although seemingly contradictory, when mitochondria are defective, despite decreased capacity to perform OxPhos, total energetic output has shown to be increased^9,12^. It has been suggested that hypermetabolism drives mitochondrial disease pathophysiology due to the existence of energetic constraints which are imposed on living things^12^. Here, we use real-time extracellular flux analysis to quantify how acute norepinephrine stimulation reshapes ATP production from OxPhos and glycolysis across a range of metabolic conditions and in the context of a genetic OxPhos deficiency in human fibroblasts.

## Results

### Norepinephrine drives a dose-dependent increase in total energetic cost

To determine the bioenergetic impact of adrenergic signaling at the cellular level, we first sought to confirm the breadth of adrenergic receptor (AR) expression in primary human fibroblasts. Using bulk RNAseq data from the Cellular Lifespan Study^9^, we averaged each cell line’s gene expression over the course of its replicative lifespan, and calculated the average of four healthy control’s gene expression. We found that these cells expressed multiple receptor subtypes, with AR-α2A (ADRA2A) exhibiting the highest expressed gene out of all the other adrenergic receptors (**Figure 1A**).

**Figure 1.**
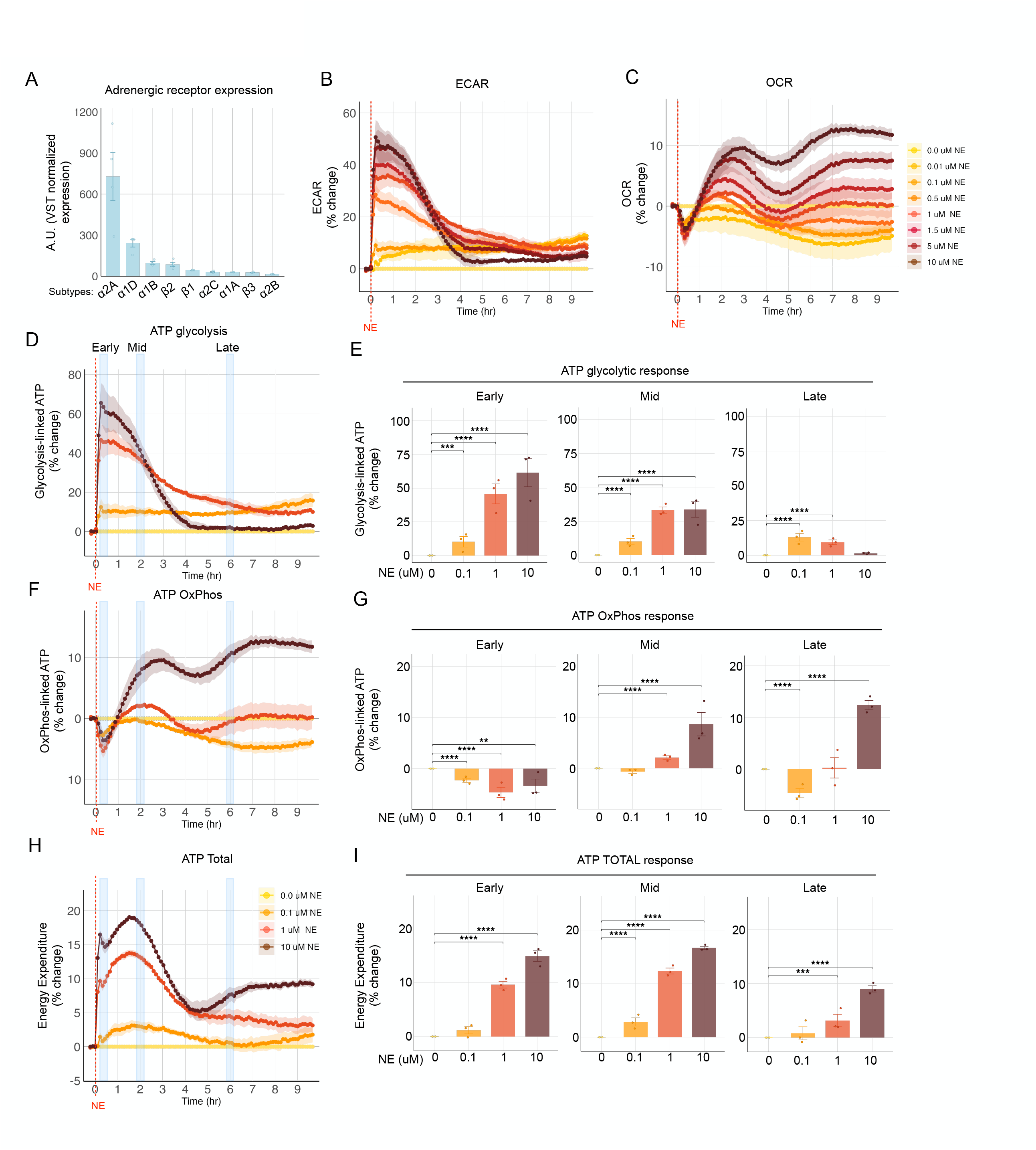
Primary human fibroblasts respond to norepinephrine in a time and dose dependent manner. (**A**) Bulk RNA expression of adrenergic receptors arbitrary units (AU, VST-normalized expression). Each receptor expression is the average of three fibroblasts lines (from independent donors), averaged across their replicative lifespan. (**B,C**) Average of three 10 hour time course experiments of fibroblasts (**B**) extracellular acidification rate (ECAR) or (**C**) oxygen consumption rate (OCR) in response to increasing doses of NE (0.0 to 10 μM NE). (**D-F**) Energy expenditure from (**D**) glycolysis, (**E**) OxPhos, or (**F**) total at select concentrations of NE. The early (18 minutes post NE injection), mid (2 hours post NE injection), and late (6 hours post NE injection) response of energy expenditure (ATP) are depicted to the right of each time course. Wells per condition per seahorse experiment n = 8-12, 20K cells per well, individually run experiments n = 3. Error bars represent SEM. Pairwise comparisons between each NE dose and vehicle control were performed using linear mixed-effects models corrected for multiple comparisons using Bonferroni method \**p* < 0.05, \*\**p* < 0.01, \*\*\**p* < 0.001, \*\*\*\**p* < 0.0001

We next stimulated fibroblasts with eight doses of norepinephrine up to 10 μM, which represents the upper end of estimated NE concentrations at neuroeffector junctions^13^ and in the ovarian parenchyma^14^. We measured OCR and ECAR at ∼6-minute intervals for up to 10 hours after NE stimulation. Norepinephrine generated a rapid increase in ECAR over the first 18 minutes post injection, with 1 μM eliciting a 34.4% increase in ECAR and 10 μM eliciting 46.7% increase in ECAR, with subsequent plateauing and eventual decline towards baseline for the remaining hours (**Figure 1B**). In contrast, OCR showed an initial decrease of -3-5% within the first 18 minutes, followed by a dose-dependent biphasic elevation, with peaks at approximately 2-2.5 hours and 7.5 hours post injection (**Figure 1C**).

Using a combination of methods for estimating ATP^15,16^, we were able to roughly estimate ATP as derived by the ECAR and OCR measurements. These calculations show that in fibroblasts, NE increases ATP from glycolysis, peaking at 9.6% (p=0.0008), 44.5% (p<0.0001), and 59.8% (p<0.0001) at early timepoints (∼18 to 36 minutes) at doses 0.1, 1, and 10 μM NE respectively (**Figure 1D**). Conversely, ATP from OxPhos dips initially to -2.3% (p<0.0001), -4.7% (p<0.0001), and -3.4% (p=0.002) at the early timepoints at those same doses, followed by a dose-dependent biphasic increase, with peaks at around 2-2.5 hours and 7.5 hours post injection of 2.1% and 8.5% (both p<0.0001) or 0.3% (ns) and 12.4% (p<0.0001) for 1 and 10 μM respectively (**Figure 1E**). In terms of total ATP (from glycolysis and OxPhos), NE caused total ATP to increase over the course of 10 hours, experiencing time and dose dependent dynamics, reaching early peaks of ATP of 1.2% (ns), 9.6% (p<0.0001) , and 15.0% (p<0.0001) over control at doses 0.1, 1, and 10 μM respectively as seen in **Figure 1F**.

### Glycolysis drives the early energetic response to norepinephrine

The rapid dries in ECAR following NE stimulation in Figure 1B Next, we sought to probe the cause for the early rise in ECAR in **Figure 1B** was suggestive that glycolysis is the primary driver of the initial energetic response. To test this directly, we employed two complementary strategies: the depletion of extracellular glucose for 24 hours prior to NE stimulation, and two by pharmacological inhibition of OxPhos via the complex V inhibitor oligomycin (Oligo) and the complex I inhibitor Piericidin A (PA). We had a fourth condition of continuous glucose (CG) as the control, and applied the physiological dose of 1 μM NE to each condition, analyzing the early response (∼18 minutes post injection).

Under all conditions, NE produced a consistent increase in glycolytic ATP: CG increase by 23.3%, no glucose by 19.8%, Oligo by 19.3%, and PA by 17.5%(**Figure 2A,D**). However, the absolute magnitude of the glycolytic response differed depending on substrate availability. In the CG condition, the increase corresponded to 55.2 pmol ATP/min, which accounted for 95% of the total increase in energy expenditure (**Figure 2G**). The no glucose condition in comparison saw a rise of 11.3 pmol ATP/min, accounting for just 28.2% of total energetic response to NE. Thus, when glucose is present, there is an approximately 5 fold greater increase in ATP production rate than without it. Interestingly, the OxPhos inhibited conditions saw an even greater ATP production rate increase of 90.5 and 77.6 pmol ATP/min for Oligo and PA respectively. Not only did these two conditions have higher baseline ATP from glycolysis, but their glycolytic energetic response was 63.8 and 40.4% greater than that of the CG condition in Oligo and PA respectively.

**Figure 2.**
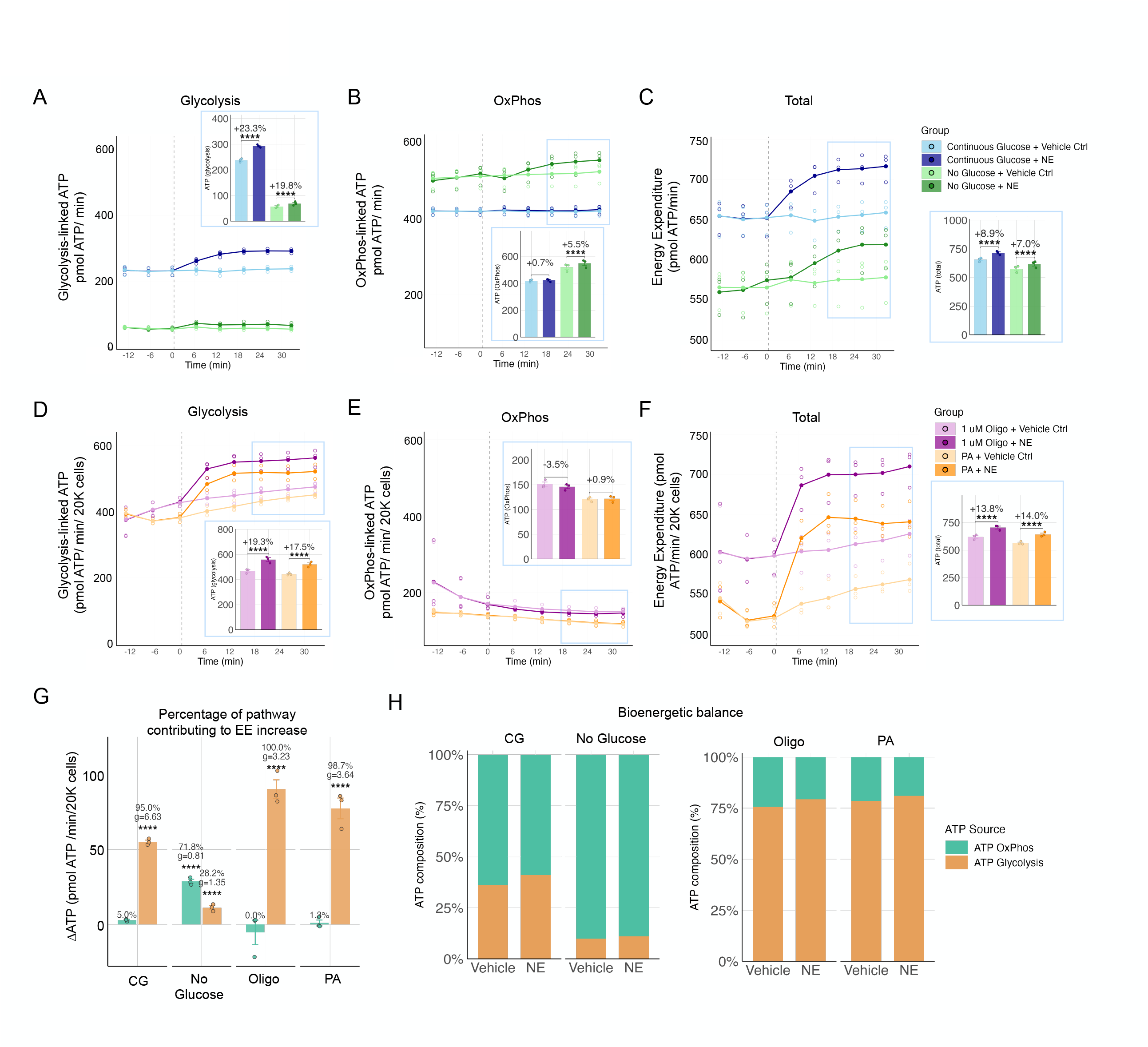
The early rise in ECAR is via glycolysis. (**A**) ATP from glycolysis in conditions of either continuous glucose or glucose depletion, ± NE. (**B**) ATP from OxPhos in conditions of either continuous glucose or glucose depletion, ± NE. (**C**) Total ATP in conditions of either continuous glucose or glucose depletion, ± NE. (**D**) ATP from glycolysis in conditions of OxPhos inhibition via PA or Oligo ± NE (**E**) ATP from OxPhos in conditions of OxPhos inhibition via PA or Oligo ± NE (**F**) Total ATP in PA or Oligo treated cells ± NE (G) ATP composition of each of the 4 conditions in A-F ∼18-36 mins post NE application. Mannitol was used instead of glucose in the injection media so as to not add glucose into the no glucose condition. (**A-F**) Each datapoint in the bar plots represent the average of the three timepoints per experiment featured in the blue outline of the scatter plots. (**G**) Percentage of each pathway contributing to the increase in ATP in response to NE for each condition. (**H**) Relative proportions of ATP from OxPhos or glycolysis per condition Continuous glucose (CG), oligomycin (Oligo), piericidin A (PA). Wells per condition per seahorse experiment n = 10-12, 20K cells per well, individually run experiments n = 3. Error bars represent SEM. Pairwise comparisons between NE and vehicle control within each metabolic condition were performed using linear mixed-effects models, corrected for multiple comparisons using Bonferroni method \**p* < 0.05, \*\**p* < 0.01, \*\*\**p* < 0.001, \*\*\*\**p* < 0.0001

OxPhos on the other had saw greatly different dynamics, in which the CG there was minimal change, an increase 2.9 pmol ATP/min or 5% of ATP from OxPhos, whereas the no glucose condition saw an increase of 28.8 pmol ATP/min which accounted for 71.8% of the increase in total ATP from OxPhos. This was 9.9x greater than the increase from the CG condition. Both the Oligo and PA conditions saw similarly trajectories, in which there was minimal to no increase in OxPhos, and all or almost all the increase came from glycolysis (100% or 98.7% respectively), as seen in **Figure 2G**.

These experiments established two main findings. One, that glycolysis was the predominant driver of the early rise in energy expenditure in response to NE when glucose is available. Second, that fibroblasts are metabolically flexible, as demonstrated when the cells were able to redirect their NE-stimulated ATP production through an alternative metabolic pathway (OxPhos). Ultimately, they further demonstrated that NE stimulated an increase in energy expenditure, regardless of pathways used.

### OxPhos-deficient SURF1-mutant fibroblasts exhibit a blunted response to norepinephrine

While pharmacological inhibition of mitochondria is indeed a method to decrease OxPhos, it is neither physiological nor is it free from having other off target effects that may confound the results of the response to NE. We therefore decided to test the energetic response to NE on patient-derived fibroblast harboring mutations in SURF1, which impairs Complex IV assembly, and ultimately reduces OxPhos capacity^9^.

Consistent with previous reports^9^, SURF1-mutant fibroblasts exhibited hypermetabolism at baseline, with total energy expenditure 11.4% greater in these cells than in healthy controls despite being deficient in OxPhos (**Figure 3C**). This excess energy expenditure was attributable to glycolysis, with SURF1-mutant cells producing 83.3% greater pmol ATP/min from glycolysis compared to the healthy controls, whereas the healthy controls had 88.7% greater pmol ATP/min from OxPhos than the SURF1-mutant cells (**Figure 3A,B**).

**Figure 3.**
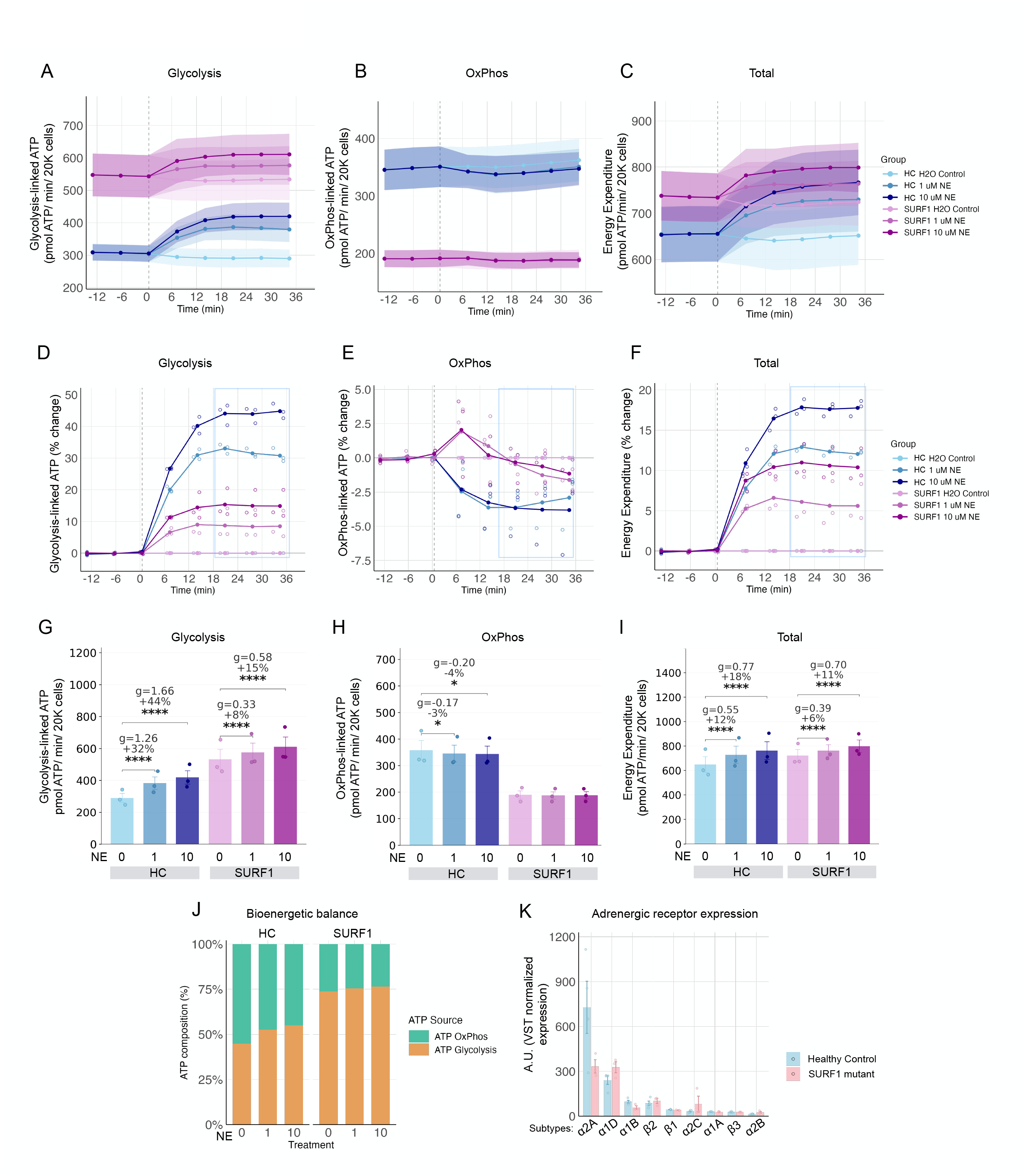
SURF1-mutant cells exhibit a blunted response to NE compared to controls. (**A**) Glycolysis-linked ATP, (**B**) OxPhos-linked ATP, and (**C**) Total energy expenditure in healthy controls and SURF1-mutant fibroblasts stimulated with 0, 1 or 10 μM NE (**D**) Percent change in glycolysis-linked ATP energy compared to controls from **A**. (**E**) Percent change in OxPhos-linked ATP compared to controls from **B**. (**F**) Total percent change in energy expenditure from **C**. (**G-I**) Energy expenditure (ATP) ± SEM from (**D-F**) at timepoints ∼18-36 mins post NE application, light blue rectangle denotes timepoints (**J**) ATP composition of each of the 4 conditions in **G-I** ∼18-36 mins post NE application. (**K**) RNAseq expression (VST normalized expression) of adrenergic receptors of healthy control cell lines (n=4) and SURF1-mutant cell lines (n=3) averaged across their lifespan. Wells per condition per seahorse experiment n = 6-10, 20K cells per well, individually run experiments n = 3. HC: healthy control; SURF1: SURF1-mutant cells. Error bars represent SEM. Pairwise comparisons between each NE dose and H_2_O vehicle control within each genotype were performed using linear mixed-effects models, corrected for multiple comparisons using Bonferroni method \**p* < 0.05, \*\**p* < 0.01, \*\*\**p* < 0.001, \*\*\*\**p* < 0.0001

When stimulated with NE, SURF1-mutant cells mounted a measurable yet blunted energetic response compared to healthy control cells. At 1 and 10 μM NE, healthy control fibroblasts increased glycolytic ATP production by 92.6 and 128.7 pmol ATP/min (31.9% or 44.3%) respectively. The SURF1-mutant cells increased by only 43.3 and 77.9 pmol ATP/min, or by 8.1% and 14.6% respectively (**Figure 3D,G**). The healthy control cells increased their ATP production from glycolysis by 2.1 and 1.7 fold compared to SURF1-mutant cells at 1 and 10 μM NE (**Figure 3I**). This is in contrast to the pharmacological inhibition in which the OxPhos inhibited cells saw the greater glycolytic response (**Figure 2G**). As expected, there was no significant change in OxPhos in the SURF1-mutant cells, while the healthy control cells saw similar initial decrease as in **Figure 1F (Figure 3B,E,H)**. Ultimately, while the SURF1-mutant cells did increase their energy expenditure in response to NE, the healthy control cells increased their total pmol ATP/min by 95.6% at 1 μM NE, and by 49.6% at 10 μM NE (**Figure 3I**).

Ultimately, the SURF1-mutant cells exhibited a more glycolytically dominant bioenergetic profile compared to healthy controls (**Figure 3J**), in line with prior findings^9^. To explore whether differences in adrenergic receptor expression could contribute to the observed genotype differences in NE responsiveness, we compared receptor subtype expression between the healthy and SURF1-mutant cells. The most highly expressed subtype, AR-α2A, was 54.3% lower in the SURF1-mutant cells compared to the healthy control cells (**Figure 3K**).

## Discussion

This study provides a systematic characterization of the bioenergetic response to norepinephrine in primary human fibroblasts, revealing that adrenergic signaling increases total cellular energy expenditure in a dose and time dependent manner. Three main observations emerge from these data.

First, NE produced a sigmoidal dose-dependent increase in total energy expenditure across 7 doses over a 10-hour time period. The temporal dynamics of this response was characterized by an early glycolytic rise, peaking at around 18 minutes, followed by a gradual decline back towards baseline, with a concomitant biphasic increase in OxPhos, first peaking at around 2 hours.

Second, fibroblasts demonstrated metabolic flexibility in in their response to NE. When glucose was depleted, cells compensated by increasing OxPhos to meet the energetic demands imposed by adrenergic signaling. Conversely, when OxPhos was inhibited pharmacologically, glycolysis alone sustained the energetic response. This is suggestive of adrenergic signaling the cells to increase energy expenditure operates upstream of some metabolic branch point, with the specific energy transforming pathway being engaged may be determined by substrate availability or enzymatic / complex capacity rather than the downstream signaling cascade of these GPCRs alone.

Third, fibroblasts from patients with SURF1 mutations exhibited a blunted energetic response to NE. These cells, which operate in a hypermetabolic state at baseline^9^, showed approximately half the glycolytic response of health controls, and as expected, lacked dynamics in OxPhos. The healthy control group increased their total energy expenditure by 12.4-17.6% while the OxPhos-deficient SURF1-mutant cells showed an increase of only 5.7-10.6%. Interestingly, despite these differences in effect size, the healthy control cells 10 μM NE condition appears to be converging on that of the SURF1-mutant cells energy expenditure, as it is only 4.7% less compared to the baseline difference of 11.4%. This could be cellular model of the constrained energy expenditure model postulated by Ponzter et al^17^, which had seen a plateauing of total energy expenditure with increasing physical activity in a diverse sample of adults. The convergence of energy expenditure at higher doses of NE is suggestive that the fibroblasts shared an upper boundary of cellular energy expenditure.

There are a few limitations to take note concerning this work. One, although it was not necessary to address the primary question of this study – the energetic cost of the adrenergic stress response – the specific adrenergic receptors that mediates the response was not investigated. Considering the difference in expression of the subtype AR-α2A between the healthy control cells and the SURF1-mutant cells, this receptor would be a strong candidate for the stress reactivity displayed throughout this work. Two, 24 hours may not have been sufficient to completely deplete the cells of all of their glycogen stores. This means for the “no glucose” condition, there may have been some residual glycogen stores and thus a possible existence of glucose present in the cells to use as a substrate for glycolysis. It is possible that with either a longer glucose depletion or the addition of a glycolytic inhibitors, such as 2-deoxyglucose, that NE would no longer produce a significant increase in ECAR or from glycolysis. Ultimately, though the rise was significant, the relative amount of ATP generated in this condition from glycolysis accounted for only 28.2% of the increase compared to 95% in the condition with glucose (**Figure 2G)**. Last, this work was done in one healthy control and one SURF1-mutant fibroblast line. Future experiments with more donor cell lines should be performed to confirm the results from this study.

Taken together, these findings position adrenergic stimulation as an energetically demanding signal, upstream of the metabolic pathway utilized to meet the demand. The blunted response of the OxPhos-deficient SURF1-mutant fibroblasts also offers insight into mitochondrial disease physiology. If cells with compromised mitochondria generate a smaller absolute increase in ATP per unit of adrenergic signal, then achieving a comparable energetic response would require greater sympathetic stimulation.

## Methods

### Fibroblast Lifespan Study

The lifespan study is explained in full detail in Sturm et al^11^. RNAseq data was downloaded from the shiny app URL: https://columbia-picard.shinyapps.io/shinyapp-Lifespan_Study/. A total of 339 samples (excluding control line technical replicate data) was used for further analysis. Five healthy control lines and three patient-derived SURF1-mutant cell lines were cultured, with or without various metabolic treatments, as described in Sturm et al.^9,11^.

### Tissue Culture

Two fibroblast lines were used: one obtained from LifeLine Cell Technology (cat # FC-0024 Lot # 00967), named hFB13 in-house, as well as the SURF1-mutant cell line received from the Hirano Lab, hFB8^11^. Fibroblasts were cultured at 5% CO_2_ and atmospheric (∼21%) O_2_. Media was low glucose (1 g/L) containing DMEM (Invitrogen #10567022), 10% FBS (Life Technologies #10437036), 50 μg/ml uridine (Sigma-Aldrich #U6381), 1% MEM non-essential amino acids (Life Technologies #11140050) and 10 μM BSA-palmitate saturated fatty acid complex (Caymon #29558). Routine passaging was performed every 5 days (± 2 days).

### Seahorse Experiments

Seahorse XF DMEM Medium, pH 7.4 with 5 mM HEPEs without phenol red, sodium bicarbonate, glucose, L-glutamine and sodium pyruvate (#103575-100) was used as the base media. Media was supplemented with 50 μg/ml uridine, 10 μM BSA-palmitate saturated fatty acid complex, 5.5 mM glucose (Thermofisher #15023021), 1 mM pyruvate (Sigma-Aldrich #S8636-100ML), 1 mM GlutaMAX Supplement (Thermo Scientific #35050061).

Cells were seeded onto Seahorse XFe96/XF Pro Cell Culture Microplates (Agilent #193794-100) at 20K cells per well (excluding the four corner wells to be used as background) the day before experiment start. 10-12 wells were seeded per group per experiment, and experiments for Figure 2 and Figure 3 used well layouts that allowed for unbiased measurements. Three independent experiments were performed for each condition. The XF sensor cartridge was hydrated with H_2_O overnight, and ∼1 hour prior to assay start was exchanged for XF calibrant solution (Agilent #100840-000). The supplemented Seahorse media was exchanged ∼1 hour prior to the start of the assay, incubated in a non-CO_2_ incubator at 37ºC. L-(-)-Norepinephrine (+)-bitartrate salt monohydrate (NE) (Sigma-Aldrich #A9512-250MG) was reconstituted to 100 mM in sterile H_2_O. Final concentrations for experiments ranged from 0.01 to 10 μM NE in the well. The mitochondrial inhibitors oligomycin (Sigma-Aldrich #75351) and piericidn A (Cayman Chemical Company #15379) were both used at a final concentration of 1 μM in the wells.

Each measurement consisted of 3 minutes of mixing following by 3 minutes of measuring, for a total ∼6 minute intervals. Experiments had either 2-6 hours of baseline measurements, then NE was injected, followed by 6-10 hours of measurements. The following day, Hoechst 33342 (Thermo Fisher Scientific #62249) was added to the wells at a final concentration of 2 μM, then counted the nuclei using the Cytation1 instrument to normalize OCR and ECAR to cell count per well.

Cell counts were run on the Cytation1 after the experiment to be able to normal to cell number. Wells with cell counts falling outside of 1.5 times the interquartile range were excluded as outliers. Cell counts across the three experiments were used to generate a grand-mean plate normalization factor to account for plate-to-plate variability. Specifically, OCR and ECAR values were normalized to cell number by dividing each well’s rate by the ratio of its cell count to the grand mean cell count computed across all non-outlier wells from all three independent experiments, yielding rates expressed at a standardized cell density. OCR and ECAR was baseline corrected by subtracting the average of the three pre-injection measurements from all subsequent timepoints such that t = 0 corresponded to the last baseline measurement before injection.

For the set of experiments investigating glucose contribution to ECAR, the media for all NE injections used mannitol (Sigma-Aldrich #M4125) to ensure osmolarity remained comparable between conditions. The condition containing no glucose were seeded with media containing mannitol instead of glucose the day of cell plating (24 hours prior to analyzing on the Seahorse). For experiments in **Figure 2**, DMSO was used as the vehicle control across all conditions to ensure a consistency of the control groups. For **Figure 1** and **Figure 3**, water was used as the vehicle control.

### ATP Calculations

ATP production rates from oxidative phosphorylation and glycolysis were estimated from the baseline-corrected OCR and ECAR using a pseudo-ATP calculation framework adapted from Desousa et al.^16^. Due to not performing the Mito Stress Test, proton leak from OCR is assumed to be 0, though is likely higher.

First, the total proton production rate (PPR_total_) was calculated from the baseline-corrected ECAR (ECAR_bc_):

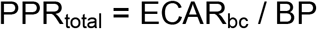

Where BP is the empirically derived buffering power (0.0909 mPH/pmol H+) which was determined by following the Agilent Seahorse XF Buffer Factor Protocol, where known amounts of acid (HCl) were injected into the assay medium. The respiratory contribution to proton production rate (PPR_resp_) was estimated using the CO_2_ contribution factor (CCF = 0.38) generated by Desousa:

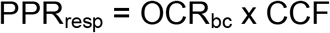

They glycolytic proton production rate (PPR_glyco_) was obtained by subtracting the respiratory contribution to PPR_total_:

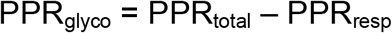

ATP production from OxPhos (ATP_OxPhos_) was calculated using the phosphate-to-oxygen ration (P/O = 2.727) calculated by Desousa et al.

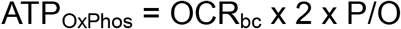

Where the factor of 2 accounts for the two atoms of oxygen per molecule of O_2_ consume. Glycoltyic ATP production (ATP_glcyo_) was calculated from the glycolytic proton production rate using the ATP-to-lactate stoichiometry (1.53 ATP per lactate), also define by Desousa et al.

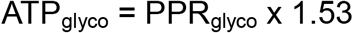

Total ATP production rate (ATP_total_) was defined as the sum of ATP_OxPhos_ and ATP_glyco_

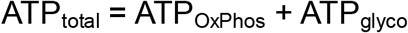

### Statistics

All analysis were performed in R (v4.4.1). Data processing, normalization, and ATP calculations were performed using R.

For each metric (OCR, ECAR, ATPOxPhos, ATPglyco, ATPtotal), measurements were analyzed within three temporal windows post-injection: Early (∼18-36 minutes), Mid (∼111-129 minutes), Late (∼471-489 minutes). Each window consisted of 3 measurements, across the 3 experiments, consisting of 9 per group (3 measurements x 3 experiments). The per-experiment mean values were averaged across technical replicate wells (ranging from n=6-12), generating one value per experiment.

Treatment effects of NE were assessed using a linear mixed-effects model. Multiple comparison correction was applied across all pairwise tests within an experiment (all temporal windows x all metrics x all pairs) using the Bonferroni method. Corrected p-values below =0.05 were considered statistically significant, with levels annotated on figures as * p < 0.05, ** p < 0.01, *** p < 0.001, **** p < 0.0001. Effect sizes were quantified as Hedge’s g^18^, as well as percent change.

## Acknowledgements

We thank Dr. Hirano for supplying the patient-derived SURF1-mutant fibroblast lines used in this study.

## Author contributions

J.L.M.S. and M.P. conceived the project. J.L.M.S. performed the experiments and analyses. J.L.M.S. generated the figures. J.L.M.S., G.S., and M.P. reviewed and edited the final manuscript.

## Funding

This work was supported by NIH grants R35GM119793, R01AG066828, R01AG086764, the Wharton Fund, and Baszucki Group to M.P.

## Data

The Lifespan Study fibroblast dataset can be downloaded at https://columbia-picard.shinyapps.io/shinyapp-Lifespan_Study/. R code for the analyses will be made available at www.github.com/mitopsychobio

## Competing Interests

The authors declare no competing interests related to this work.

## Supplementary Material

This article contains 3 Supplementary Figures.

## Notes

https://columbia-picard.shinyapps.io/shinyapp-Lifespan_Study/

### Competing Interest Statement

The authors have declared no competing interest.

