## Supplemental Figure 1 for "The Energetic Cost of Adrenergic Signaling in Primary Human Fibroblasts"

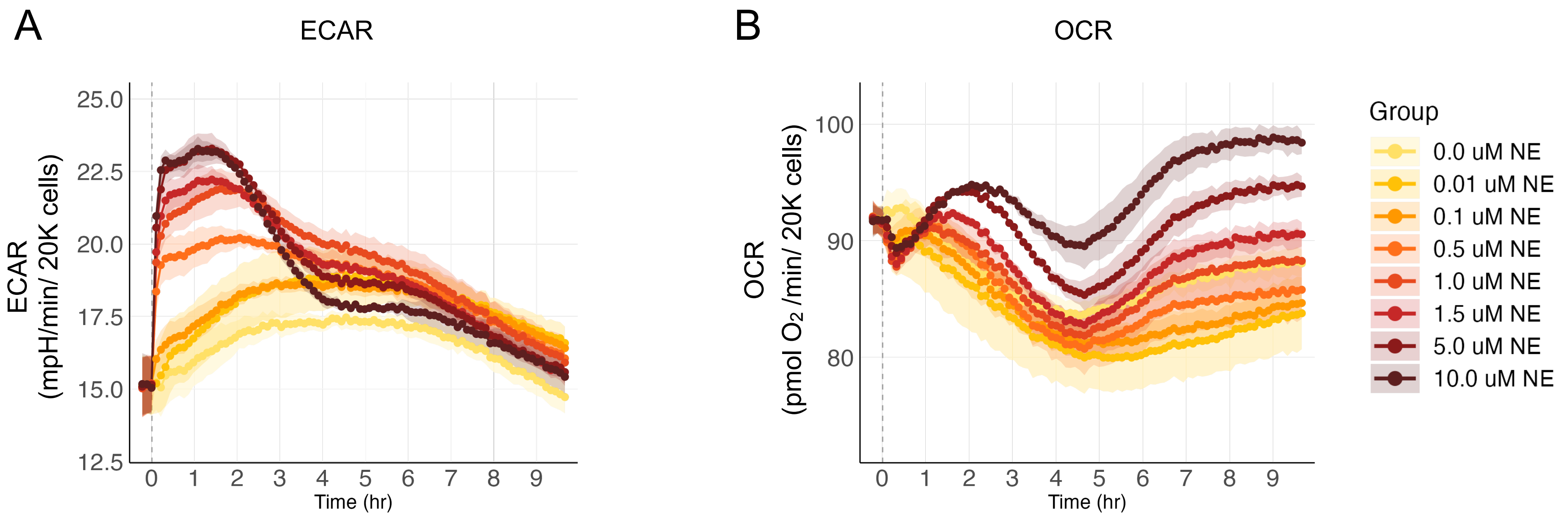


**Supplemental Figure 1. Raw values of ECAR and OCR to NE doses over 10 hours** (**A,B**) Raw average baseline-corrected values of ECAR and OCR of three separate 10 hour time-course experiments of fibroblasts in response to increasing doses of NE (0.0 to 10 μM NE). (**A**) extracellular acidification rate (ECAR) or (**B**) oxygen consumption rate (OCR). Vertical dashed line represents timing of NE or vehicle control injection. Wells per condition per seahorse experiment n = 8-12, 20K cells per well, individually run experiments n = 3.
