## Supplemental Figure 2 for "The Energetic Cost of Adrenergic Signaling in Primary Human Fibroblasts"

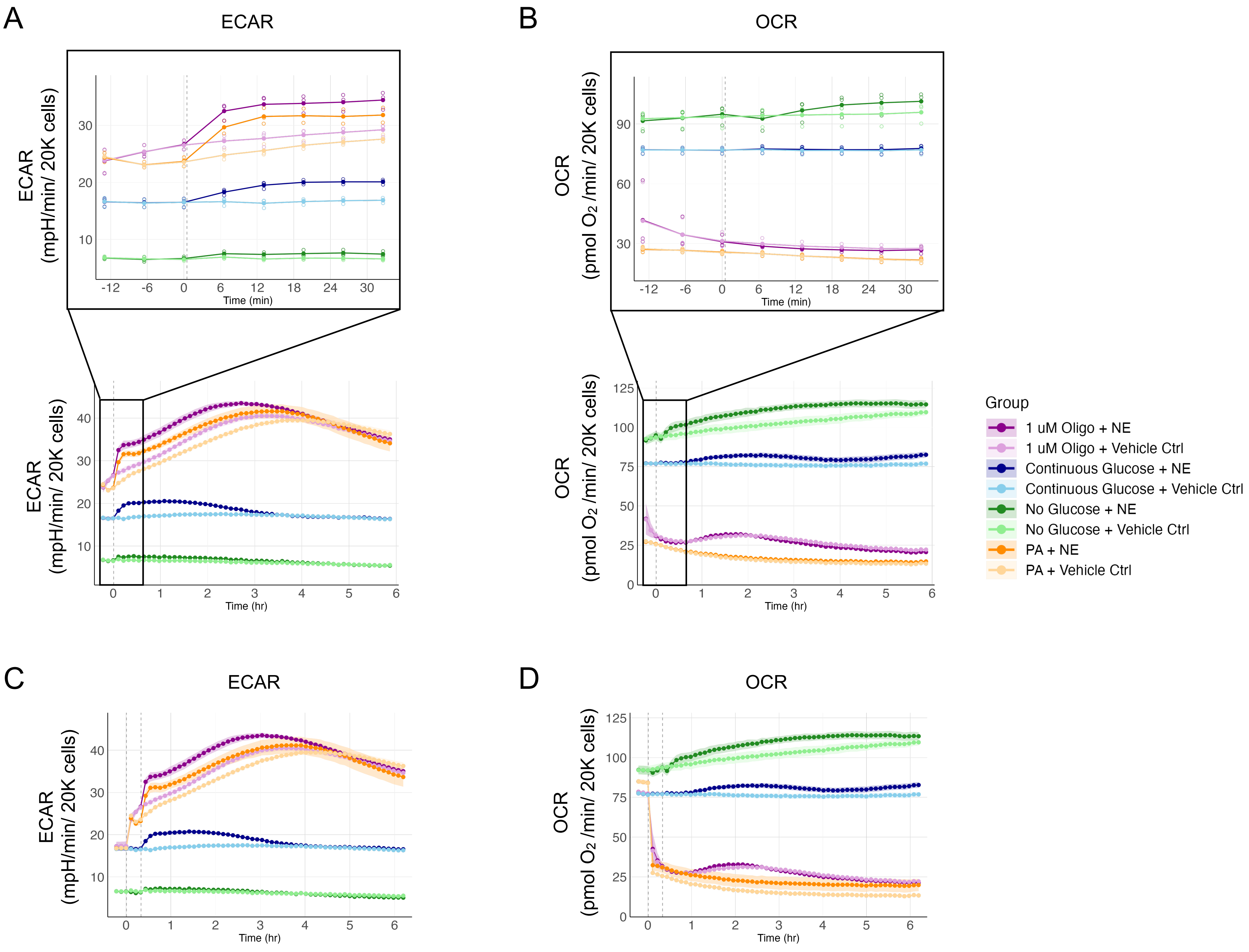


**Supplemental Figure 2. Raw values for mitochondrial inhibition and glucose withdrawal experiments** (**A**) Time-course extending to 6 hours post NE injection of all conditions: continuous glucose, glucose depletion, oligomycin (Oligo) or piericidin A (PA) ± NE, as well as zoomed in on early timepoints (inset). Baseline-corrected to the three timepoints before the NE injection (**A**) Extracellular acidification rate (ECAR) (**B**) Oxygen consumption rate (OCR) (**C,D**) Extended time-course depicting two injections, represented by vertical dashed lines, of either the mitochondrial inhibitors (Oligo or PA) or vehicle control, followed by the second injection of NE or vehicle control, up to 6 hours post NE injection. Baseline-corrected to the average of three timepoints before the first injection. (**C**) ECAR (**D**) OCR. Mannitol was used instead of glucose in the injection media so as to not add glucose into the no glucose condition. Wells per condition per seahorse experiment n = 10-12, 20K cells per well, individually run experiments n = 3.
