## Supplemental Figure 3 for "The Energetic Cost of Adrenergic Signaling in Primary Human Fibroblasts"

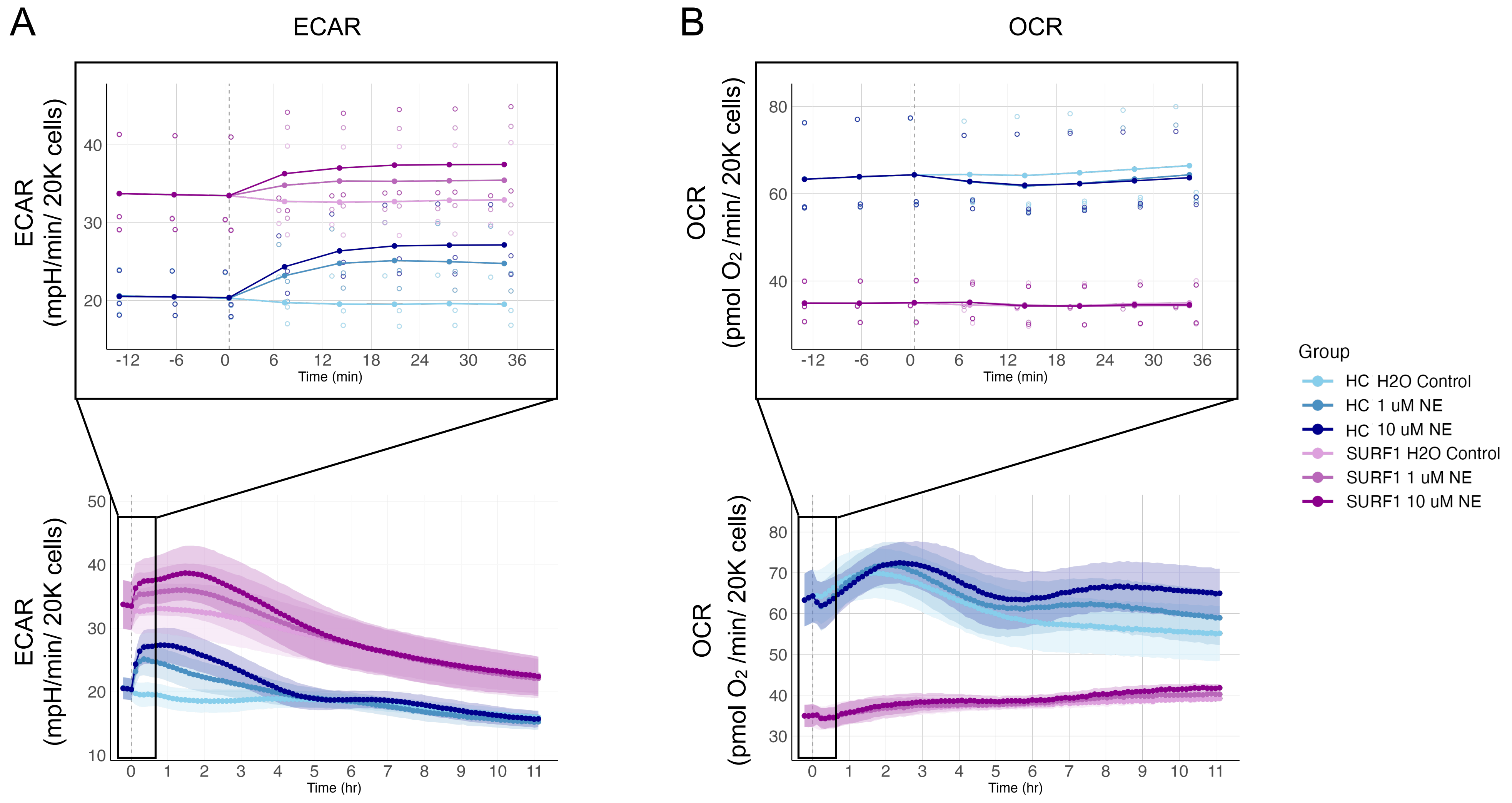


**Supplemental Figure 3. Raw values for healthy control and SURF1 mutant time-courses** (**A**) Time-course extending to 6 hours post NE injection of healthy control (HC) cells vs SURF1 mutant cells ± NE, as well as zoomed in on early timepoints (inset). Baseline-corrected to the three timepoints before the NE injection (**A**) Extracellular acidification rate (ECAR) (**B**) Oxygen consumption rate (OCR). Vertical dashed line represents timing of NE or vehicle control injection. Wells per condition per seahorse experiment n = 6-10, 20K cells per well, individually run experiments n = 3.
